# Bacterial toxin-antitoxin system MazEF as a native defense mechanism against RNA phages in *Escherichia coli*

**DOI:** 10.1101/2023.02.01.526697

**Authors:** Nela Nikolic, Tobias Bergmiller, Maroš Pleška, Călin C. Guet

## Abstract

Bacteria have evolved a wide range of defense strategies to protect themselves against bacterial viruses (phages). However, the known mechanisms almost exclusively target phages with DNA genomes. While several bacterial toxin-antitoxin systems have been considered to cleave single-stranded bacterial RNA in response to stressful conditions, their role in protecting bacteria against phages with single-stranded RNA genomes has not been studied. Here we investigate the role of a representative toxin-antitoxin system, MazEF, in protecting *Escherichia coli* against two RNA phages – MS2 and Qβ. Our population-level experiments revealed that a *mazEF* deletion strain is more susceptible to RNA phage infection than the wild-type. At the single-cell level, deletion of the *mazEF* locus significantly shortened the time to lysis of individual bacteria challenged with RNA phage. At the genomic level, we found that the adenine-cytosine-adenine sequence, directly recognized and cleaved by the MazF toxin, is systematically underrepresented in the genomes of RNA phages that are known to infect *E. coli*, indicating selection for decreased probability of cleavage. These results suggest that in addition to other physiological roles, RNA-degrading toxin-antitoxin modules can function as a primitive immune system against RNA phages.

## INTRODUCTION

Phages are widely abundant viruses that infect Bacteria and Archaea. Their genomes can be composed of single- or double-stranded DNA or RNA. Bacteria have evolved different defense mechanisms to protect themselves against phages. So far, almost all studies that investigate bacteria fighting phage infection have addressed defense systems against phages with DNA genomes [Labrie et al 2010, Samson et al 2013, Bernheim and Sorek 2020, Vassallo et al 2022], raising the question of how bacteria defend themselves against phages with RNA genomes. MS2 (*Emesvirus zinderi*) and Qβ (*Qubevirus durum*) are the best-known bacteriophages with genomes composed of a positive-sense single-stranded RNA (or (+)ssRNA), which means that the genome also directly serves as mRNA that is translated into proteins inside the host cell [van Duin and Tsareva 2006]. (+)ssRNA bacteriophages (referred to as RNA phages hereafter) start infection by adsorbing to bacterial pili, and induce lysis of infected cells.

The main bacterial defense strategies against phages with single-stranded and double-stranded DNA genomes that were discovered so far, include cell surface modifications to avoid phage adsorption, cleavage of viral nucleic acids by restriction-modification or CRISPR-Cas systems, and induction of cell suicide to abort phage infection through various Abi systems [Samson et al 2013, van Houte et al 2016, Rostøl and Marraffini 2019, Negri et al 2021]. However, many underlying mechanisms and their influence on bacterial physiology still remain elusive, with recent studies indicating that there are many *unknown* antiphage systems for protection against *well-known* phages [Doron et al 2018, Barrangou and van der Oost 2018, Vassallo et al 2022]. Moreover, even for defense systems that have been known for a long time, we still lack a basic understanding of how the underlying molecular mechanisms impact bacteria-phage interactions at the ecological scale [Pleška et al 2016, Pleška and Guet 2017, Pleška et al 2018]. For example, CRISPR-Cas systems, which are traditionally viewed as defense mechanisms against DNA phages, have been shown to recognize and cleave ssRNA [Yan et al 2019], including the RNA genome of the MS2 phage [Abudayyeh et al 2016, Smargon et al 2017, Strutt et al 2018]. Moreover, several toxin-antitoxin (TA) systems have been implicated in cleaving ssRNA in response to stressful conditions [Gerdes 2012], however their role in protecting bacteria against RNA phages has not been assessed.

Here we investigate the role of the TA system MazEF in defending *E. coli* against RNA phages. Generally, TA modules are genetically encoded systems widespread across bacterial genomes [Van Melderen 2010, Yamaguchi and Inouye 2011, Gerdes 2012, Song and Wood 2020]. Toxin activation during stressful conditions inhibits basic cellular processes, for instance by interfering with replication, translation or cell wall synthesis. Chromosomally encoded TA systems protect against invading plasmids and DNA phages, modulate bacterial gene expression and growth, and collectively help bacteria to withstand stressful conditions [Magnuson 2007, Van Melderen 2010, Nikolic et al 2017, Nikolic et al 2022]. TA systems have already been implicated as defense tools against DNA phages through different mechanisms of direct interference [Song and Wood 2020, LeRoux et al 2022]. Moreover, it has also been shown that DNA phage infection can disrupt the cellular toxin-antitoxin balance and promote toxin activation, which consequently reduces phage production [Magnuson 2007, Guegler and Laub 2021]. This raises the potential of TA systems to act as Abi systems, and mediate abortive infection by inducing bacterial cell suicide [Labrie et al 2010]. To what extent and by what means TA systems protect their host from RNA phages remains an open question.

The MazEF system is one of the most abundant TA systems across *E. coli* species, present in the genomes of over 80% of *E. coli* strains [Norton and Mulvey 2012, Fiedoruk et al 2015, Ramisetty and Santhosh 2015]. The MazF toxin recognizes an ACA sequence as the core cleavage site in single-stranded RNA of *E. coli* [Zhang et al 2003, Zorzini et al 2016, Culviner and Laub 2018]. It has been considered that MazF cleaves RNA in the presence of stress, as for instance during nutrient depletion [Mets et al 2017]. The model *E. coli* strain K-12 MG1655 has at least ten type II toxins-endoribonucleases that degrade RNA, and few of them, such as MazF, exhibit sequence-specificity [Harms et al 2018]. The RNA genome of phage MS2 has been commonly used to determine *in vitro* RNA cleavage mediated by MazF of *E. coli* [Zorzini et al 2016] and of other species [Nariya and Inouye 2008, Zhu et al 2008, Zhu et al 2009, Park et al 2011]. Even though RNA phage genomes are highly structured, experimentally determined secondary structures of RNA phages MS2 and Qβ indicate that ACA motifs can be found in unpaired segments of the loop regions, i.e. single-stranded regions [Olsthoorn et al 1996, Klovins et al 1998], and thus accessible to MazF cleavage.

While MazEF is a biochemically well-characterized TA system, its physiological roles have so far remained controversial [Jurenas et al 2022]. In this study, we investigate interactions between the MazEF system and RNA phages MS2 and Qβ. Our experimental analysis indicates that MazEF is a part of bacterial native defense machinery against RNA phage infection, further suggesting that TA systems can act as a primitive immune system against RNA phages.

## MATERIAL AND METHODS

### Strains

*E. coli* strain K-12 MG1655 [Blattner et al 1997] and its derivatives were used in this study. All bacterial strains and plasmids as well as RNA phages are listed in **Supplementary Table S1**. *E. coli* strain NN238 was made by P1-transduction of Δ*araA*::*kanR* from TB315 to LVM101 [Tsilibaris et al 2007]. Strains NN239 and NN241 were made by using the CRIM method described in [Haldimann and Wanner 2001]: the sequence frt-*cat*-frt (harboring chloramphenicol resistance gene *cat*) from modified CRIM plasmid pAH68-frt-cat was inserted into its attachment site *att*HK022. Strains NN242-cat and NN243-cat were made by P1-transduction of *att*P21::λP_R_-*mCherry*::frt-*cat* from TB194 to MG1655 LVM and LVM101, respectively. The chloramphenicol resistance marker was removed by using the site-specific Flp-recombinase [Cherepanov and Wackernagel 1995], resulting in strains NN242 and NN243. F plasmid from strain W1485 was introduced by conjugation into strains TB315, NN238, NN239, NN241, NN242-cat, and NN243-cat.

### Bacterial cultivation

Cultures were grown in rich LB medium, defined rich or minimal medium. Defined rich medium contains 1× M9 salts, 1 mM MgSO_4_, 0.1 mM CaCl_2_, 0.5% casamino acids, 10 mM maltose. Minimal medium contains 1× M9 salts, 1 mM MgSO_4_, 0.1 mM CaCl_2_, 30 mM maltose. Generally, frozen glycerol clones were first streaked on LB agar plates. All incubations were carried at 37°C, with batch cultures shaking at 230 rpm. Prior to adding RNA phage for infection experiments, 0.01% glucose and 2 mM CaCl_2_ were added to bacterial cultures. Where appropriate, the antibiotics were added to medium in the following final concentrations: 15 μg/ml chloramphenicol and 100 μg/ml ampicillin for plasmid maintenance, and 10 μg/ml chloramphenicol and 25 μg/ml kanamycin for selection after P1 transduction and conjugation.

### Phage lysate preparation

Overnight cultures of *E. coli* W1485 strain were diluted 1 to 100 into fresh LB medium for 4 h. Exponentially growing bacterial cultures were then inoculated with a single plaque in 4 ml of phage soft agar (1% tryptone, 0.1% yeast extract, 0.01% glucose, 0.8% NaCl, 2 mM CaCl_2_, 0.7% agar; kept at 50°C) and plated on phage plates (1% tryptone, 0.1% yeast extract, 0.01% glucose, 0.8% NaCl, 2 mM CaCl_2_, 1% agar). Plates were incubated overnight at 37°C for 20 h. Next day, soft agar containing plaques was scrapped off into a 50 ml-Falcon tube, 12 ml of SM buffer was added (100 mM NaCl, 8 mM MgSO_4_·7H_2_O, 50 mM Tris-Cl pH 7.5, 0.01% gelatin), and the Falcon tubes were centrifuged for 15 min at 4000 g. The supernatant (phage lysate) was sterilized twice through a 0.22 μm-filter, and kept in fridge at 4°C. The phage titer was assessed with plaque assays.

### Plaque spotting assays

To determine the phage titer in phage lysates, overnight cultures of host strain W1485 were diluted 1 to 1000 into fresh LB medium, and cultivated for 5.5 h. 200 μl of exponentially growing bacterial cultures were then added to 4 ml of phage soft agar, briefly vortexed, and plated on phage plates. After 2 min, 5 μl of serial dilutions of the phage lysate were spotted on the phage plates containing bacterial host. Plates were incubated overnight for 20 h. The number of plaque-forming units PFU per ml of the lysate was calculated as: PFU/ml= n_plaques_ / (dilution factor * V_diluted phage_).

### Colony count after phage infection

A single colony for the wild-type NN239 F+ or Δ*mazEF* NN241 F+ strain was inoculated in LB medium and cultivated overnight 17-19 h. The cultures were diluted 1 to 1000 into 3 ml of medium and cultivated for 3.5 h. Each exponentially growing bacterial culture was split: one part remained uninfected, while the second part was infected with RNA phage at the specific multiplicity of infection (MOI, the ratio of phage particles to bacterial cells in a culture). Samples were incubated at 37°C without shaking for 27 min for MS2 (15 min for Qβ), to allow for phage adsorption in infected samples, then incubated at 37°C with shaking for total infection time of 90 min for MS2 or 105 min for Qβ. All samples were then washed with SM buffer to wash away the non-adsorbed phage, and then re-suspended in fresh LB medium. Serial dilutions were plated on LB agar plates, then incubated for 24 h. The number of bacterial cells in mixed cultures was determined by measuring colony forming units as: CFU/ml= n_colonies_ / (dilution factor * V_diluted culture_). CFU measurements were normalized by the colony count before phage infection at *t*= 0 min.

### Plate-reader experiments

Overnight cultures of the wild-type NN239 F+ and Δ*mazEF* NN241 F+ strains were diluted 1 to 1000 into 4 ml of LB medium. After 3 h of cultivation, the cultures were supplemented with 0.01% glucose and 2 mM CaCl_2_ in the final concentrations, and 195 μl of the cultures were put into a 96-well plate. The cultures were either infected with 5 μl of RNA phage lysate at MOI= 1, or 5 μl of SM buffer was added to uninfected cultures. Growth of the cultures was recorded every 5 min for 10 h in total, as absorbance at 600 nm A_600_, with CLARIOstar Plate Reader, BMG Labtech.

### Strain labeling for competition assays

Strain NN239 F+ (*mazF*+ Ara+) was competed against strain NN238 F+ (*mazF*-Ara-). Strain TB315 F+ (*mazF*+ Ara-) was competed against strain NN241 F+ (*mazF*-Ara+). The strains that are Ara+ can metabolize arabinose, specifically, those strains carry functional *araA*. An Ara+ strain produces pinkish-white colonies, whereas an Ara-strain produces red colonies when grown on tetrazolium arabinose agar plates (1% tryptone, 0.1% yeast extract, 1% arabinose, 0.5% NaCl, 0.005% triphenyltetrazolium chloride, 1.6% agar), i.e. the red-white coloration of colonies depends on the *araA* genotype. The *araA* mutation makes a suitable neutral marker for competition experiments [Leon et al 2018].

### Competition assays in rich LB medium

Competition assays were performed to evaluate if the wild-type or Δ*mazEF* strain are better adapted to RNA phage infection. Competition pairs of exponentially growing bacterial cultures were mixed in approximately 1:1 ratio. Each mixed culture was split: One part remained uninfected, while the second part was infected with RNA phage. Exact numbers of bacteria and phage particles are indicated in figure legends. Samples were incubated at 37°C without shaking for 27 min for MS2 (15 min for Qβ), then incubated at 37°C with shaking for total infection time of 90 min for MS2 or 105 min for Qβ. All samples were then washed with SM buffer to wash away the non-adsorbed phage, and then re-suspended in fresh LB medium. Serial dilutions were plated on tetrazolium arabinose agar plates, then incubated for 24 h. The number of bacterial cells in mixed cultures was determined by measuring colony forming units of red and white colonies as CFU/ml. The Wrightian fitness of the wild-type strain NN239 F+ was determined relative to the fitness of the Δ*mazEF* strain NN238 F+ at time *t*, and normalized by the colony count before addition of phage at *t*= 0 h: relative fitness= (CFUwt_t_/CFUwt_0_) / (CFUΔ_t_/CFUΔ_0_).

### Competition assays in minimal medium

A single colony for each strain was inoculated in defined rich medium and grown for 6 h. The cultures were diluted 1 to 100 into 3 ml of minimal medium and cultivated overnight 17-19 h. On the following day, the cultures were diluted 1 to 100 into fresh minimal medium for 5 h. Competition pairs were mixed in approximately 1:1 ratio. Each mixed culture was split: One part remained uninfected, while the second part was infected with RNA phage. Exact numbers of bacteria and phage particles are indicated in figure legends. Samples were incubated without shaking for 30 min (to allow for phage adsorption in infected samples), then incubated at 37°C for 18-20 h overnight. Next day, all samples were washed with SM buffer to wash away the non-adsorbed phage, and then re-suspended. Serial dilutions were plated on tetrazolium arabinose agar plates, then incubated for 24 h.

### Colony count after *mazF* overexpression and phage infection

Overnight cultures of strain NN238 F+ pBAD-*mazF* were diluted 1 to 500 into 4 ml of defined rich medium with chloramphenicol for 2.5 h, to obtain exponentially growing bacterial populations. Each culture was then divided: One part remained uninduced, while 0.01% arabinose (Ara) was added to the second part to induce *mazF* expression. After 30 min, all cultures were again divided: One part remained uninfected, while the second part was infected with RNA phage. Samples were incubated on ice for 30 min, then incubated at 37°C for 2.5 h with shaking. All samples were washed with SM buffer, and then re-suspended. Serial dilutions were plated on LB agar plates with chloramphenicol, then incubated overnight for 18 h. The number of bacterial cells in cultures was determined next day by measuring colony forming units as CFU/ml. CFU count of each replicate culture was measured before infection (uninduced, *mazF*-induced) and after infection (control uninduced, control *mazF*-induced, infected uninduced, infected *mazF*-induced, as indicated in **Supplementary Datasets**).

### Plaque assays after *mazF* overexpression

Overnight cultures of strain NN238 F+ pBAD-*mazF* were diluted 1 to 1000 into defined rich medium with chloramphenicol for 3.5 h. Each culture was then divided: One part remained uninduced, while the second part was *mazF*-induced by adding Ara. After 30 min to 1 h, all samples were washed with LB medium, to remove Ara. 100 μl of each washed culture containing 10^7^-10^8^/ml bacterial cells were mixed with 10 μl of phage MS2 or phage λ *vir* lysate in 3 ml of phage soft agar and plated on phage plates. Plates were incubated overnight for 20 h, and PFU/ml was determined the following day.

### Microfluidic-based time-lapse fluorescence microscopy for phage infection assays of single bacterial cells

Microfluidic devices were prepared as described in detail elsewhere [Bergmiller et al 2017, Nikolic et al 2018] and operated using NE-700 syringe pumps with a flowrate of 2 ml/h. A temperature-controlled Olympus IX83 microscope was equipped with a Lumencore SpectraX light source and a custom-made autofocus system [Chait et al 2017]. Images were acquired every 5 min using a 100× 1.4 NA oil immersion objective lens and a cooled Photometrix Prime95B. To image mCherry fluorescent protein, we used the green LED (549 ± 15 nm) with an intensity of 320 mW and an exposure time of 200 ms. To image fluorescein, we used the cyan LED (475 ± 28 nm) at 180 mW and 25 ms exposure time. The emission filters were from Semrock (LP 495, BP 520/35 for GFP and LP 596, BP 641/75 for mCherry). As F-pili promote biofilm formation in *E. coli* F+ strains [May et al 2011], overnight cultures of the wild-type NN242-cat F+ or Δ*mazEF* NN243-cat F+ were mixed with 0.1% Tween to enable efficient loading of bacteria into a microfluidic device and to prevent bacteria clumping within the device. Bacteria were growing for at least 3 h at 37°C to allow for a steady-state growth in LB medium, before switching to LB medium supplemented with 0.01% glucose, 2 mM CaCl_2_ and 0.001% fluorescein, and containing phage lysate in the final concentration of 10^9^ MS2 or Qβ phage particles/ml. Fluorescein was used to determine the exact timing of the media switching and exposure of bacterial cells to RNA phage. The fixed RNA phage concentration was maintained throughout the experiment, with any phage progeny immediately being washed away with the media flow. Bacterial cells at the blunt-end of each populated growth-channel were analyzed with Fiji (ImageJ) during 650 min of phage MS2 infection or 985 min of phage Qβ infection.

### Bacterial cell length analysis

We analyzed fluorescence microscopy images of *E. coli* wild-type and Δ*mazEF* cells corresponding to time of 5 min before adding phage, and 200 min after adding RNA phage. The length of all cells in the field of view was measured in units of pixels with the Matlab-based package *Schnitzcells* [Young et al 2012]. The length was then converted to units of μm using 0.11 μm/pixel as conversion factor.

### Statistical analysis

Statistics was done in Microsoft Excel, R and Matlab. To evaluate differences in CFUs of different strains we used two-tailed two-sample heteroscedastic Student’s *t*-tests. To evaluate differences in relative fitness between infected and uninfected mixed cultures we used two-tailed paired Student’s *t*-tests. To evaluate differences in the time to lysis of single bacterial cells, as well as differences in the cell length, we used non-parametric two-sample Mann-Whitney tests (different strains, different cultivation media, or different time points). Competition assays and burst size assays were evaluated by the Matlab function *fitlm* (details in **Supplementary Methods**).

### Genomes

From the National Center for Biotechnology Information’s (NCBI) Viral Genomes Database [Brister et al 2014], https://www.ncbi.nlm.nih.gov/genome/viruses/ (March 2019), we collected 12 full-length reference genomes of viruses with (+)ssRNA genomes, which infect Bacteria (RNA phages). We also collected 2216 genomes of viruses with genomes other than (+)ssRNA, which infect Bacteria, and 1835 genomes of (+)ssRNA viruses, which do not infect Bacteria. From the NCBI Nucleotide Database https://www.ncbi.nlm.nih.gov/nucleotide/, we collected 28 RNA phage genomes from [Friedman et al 2009], 20 RNA phage genomes from [Krishnamurthy et al 2016], and additional 33 complete and partial RNA phage genomes with the length of more than 1000 nucleotides and excluding genomes obtained from experimentally evolved strains, giving a total of 81 RNA phage genomes. In addition, from those 81 phages, we excluded “unclassified” members (the NCBI Taxonomy Database https://www.ncbi.nlm.nih.gov/Taxonomy/Browser/www.tax.cgi?id=11989), resulting in a total of 57 (+)ssRNA phages that infect *Escherichia*.

### Genome analysis

Percentage of ACA sites in a genome was calculated as [n_ACAgenome_ / (length_genome_ - 2)] * 100. Percentage of expected ACA sites was calculated as [fraction(A)_genome_ * fraction(C)_genome_ * fraction(A)_genome_ * 100]. Number of expected ACA sites was calculated as [fraction(A)_genome_ * fraction(C)_genome_ * fraction(A)_genome_ * length_genome_]. The relative ACA frequency was calculated as the percentage of counted ACA sites divided by the percentage of expected ACA sites in the genome. The analysis of other toxin recognition trinucleotides – GCU, ACG, ACU – was done in the same way. **Genome shuffling**: Each ssRNA phage genome was shuffled such that the nucleotide content remained the same as in the original genome (fractions of A, C, G and U were the same in the original and shuffled genomes), and the shuffling of nucleotides was repeated 10,000 times. The analysis was done in Matlab.

## RESULTS

### Deletion of the chromosomal *mazEF* locus increases RNA phage-mediated killing

In this project we aimed to elucidate interactions between the MazEF system and RNA phages. MazF of *E. coli* is an RNA-degrading enzyme that specifically recognizes ACA sequences in the single-stranded regions of *E. coli* RNA [Zhang et al 2003], and ACA sequences in foreign RNA molecules, such as the RNA genome of phage MS2 [Zorzini et al 2016]. We experimentally determined how MazEF, the TA system native to *E. coli* K-12 MG1655, influences survival and growth of *E. coli* populations during RNA phage infection. All *E. coli* strains investigated here carry the F plasmid, as phages MS2 and Qβ start infection by binding along the side of F-pili encoded on the conjugation F plasmid [van Duin and Tsareva 2006]. It should be noted that the original *E. coli* K-12 isolate carried the F plasmid [Blattner et al 1997].

We first set out to measure the effect of MS2 and Qβ infection on the viability of *E. coli* wild-type strain K-12 MG1655 F+ and the isogenic *mazEF* deletion strain. The incubation time in these experiments was 90 minutes for MS2 and 105 minutes for Qβ, corresponding to the average length of the respective phage replication cycle [Rappaport 1965, Jenkins et al 1974, Woody and Cliver 1995, Tsukada et al 2009]. Bacteria were infected at the mid- or mid-to-late exponential phase, as the maximum production of F-pili is achieved during exponential growth in aerobic cultures [Biebricher and Dueker 1984]. Infections were carried at various levels of multiplicity of infection, MOI, which is the ratio of phage particles to bacterial cells in a culture. Under these conditions, *mazEF* deletion reduced the number of colony forming units (CFU) following the MS2 infection (MOI= 0.1) and Qβ infection (MOI= 1) 2.1 and 2.3 times, respectively (1.7 and 2.5 times at MOI= 0.01) (**Figure 1A-F**). At the timescale beyond a single infection cycle, *E. coli* wild-type cultures were less affected by RNA phage, than were the Δ*mazEF* cultures (**Figure 1G-I, Supplementary Figure S1**). Specifically, bacterial population dynamics assays in a plate-reader showed that after ten hours of MS2 (Qβ) phage infection, biomass of the *E. coli* wild-type populations was on average 22% (18%) larger than biomass of the *E. coli* Δ*mazEF* populations. These findings indicate that the presence of the MazEF system lowers bacterial host mortality upon the addition of RNA phage.

**Figure 1.**
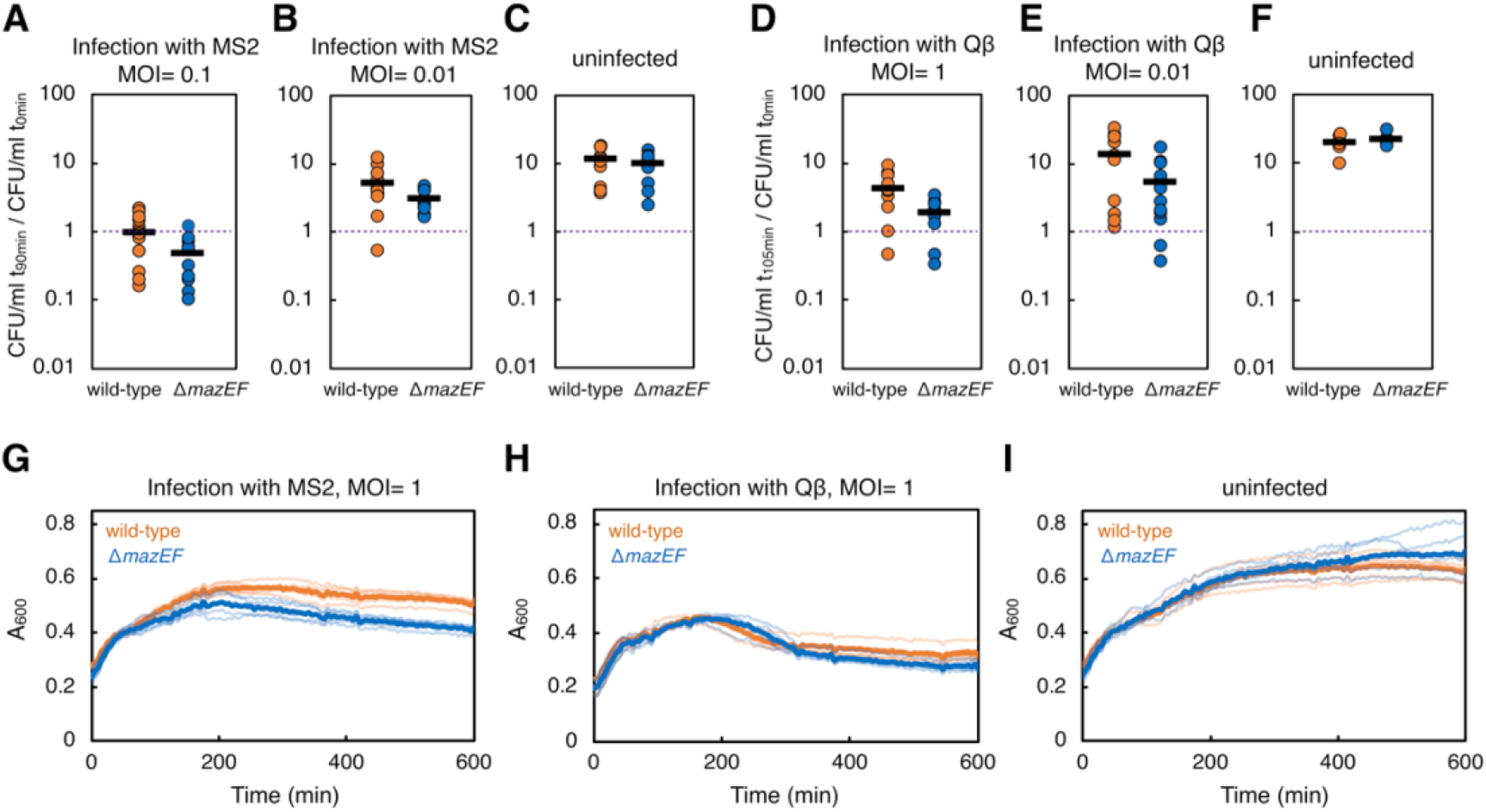
Colony count and bacterial population dynamics during exposure to RNA phage. Exponentially growing cultures of the wild-type (orange) and Δ*mazEF* (blue) strains harboring F plasmid (∼10^7^ bacterial cells) were exposed to RNA phage at below indicated multiplicity of infection, MOI. **A)** Here plotted is the density of colony forming units (CFU) at the indicated time after phage addition, normalized by the density of CFU at time 0: phage MS2, MOI= 0.1 (13 independent replicates, *t*-test *P*= 0.02), **B)** phage MS2, MOI= 0.01 (13 replicates, *P*= 0.03), **C)** uninfected cultures (12 replicates, *P*= 0.41), **D)** phage Qβ, MOI= 1 (10 replicates, *P*= 0.03), **E)** phage Qβ, MOI= 0.01 (11 replicates, *P*= 0.049), **F)** uninfected cultures (7 replicates, *P*= 0.43). **G)** Population dynamics of infected and uninfected control cultures of the wild-type (orange line) and Δ*mazEF* (blue line) strains, was recorded in a plate-reader as absorbance at 600 nm (A_600_). Phage MS2 was added to the cultures at time 0 and MOI= 1 (5 replicates, one *t*-test per each time point during period 540-600 min, 13 *t*-tests in total, all *P*< 0.00007), **H)** phage Qβ at MOI= 1 (5 replicates, *t*-tests during period 540-600 min *P*< 0.036), or **I)** no phage added (5 replicates, *t*-tests during period 540-600 min *P*> 0.185). Brighter lines represent measurements of individual cultures, darker lines are the mean values for each strain.

The presence of the *mazEF* locus did not affect RNA phage adsorption, which is the first step of phage infection cycle (**Supplementary Figure S2**). However, phage burst size assays, reporting on phage progeny production, showed that wild-type cultures produced on average 1.5 and 2.8-fold fewer plaques in a single infection step by MS2 and Qβ, respectively, compared to Δ*mazEF* cultures (**Supplementary Figure S3**). These results suggest that rather than lowering the likelihood of phage adsorption (which could occur for example due to lower expression of F-pili), MazEF interferes with RNA phage replication within already infected cells.

Significant effect of the *mazEF* deletion was also detected in a direct competition assay, in which mixed cultures of the wild-type and Δ*mazEF* strains were infected with RNA phage MS2 or Qβ for short and long timescales. Short infection was carried out for 90 minutes for MS2 or 105 minutes for Qβ in rich media, while long infection was carried out for 20 hours in minimal media. Our results showed that under both sets of conditions, the presence of *mazEF* conferred a significant advantage to *E. coli*. The relative fitness of the wild-type strain was 1.62 and 1.31 for short infection at MOI= 1 (1.29 and 1.15 at MOI= 0.1) in the case of MS2 and Qβ, respectively (**Figure 2A-D**). For long infection at MOI= 0.1, the relative fitness of the wild-type strain was 1.33 and 1.45 in the case of MS2 and Qβ, respectively (**Figure 2E-F**). In the RNA phage absence, deletion of *mazEF* did not affect the fitness significantly. Overall, the results suggest that, in the presence of RNA phage, a MazEF+ strain has a larger probability of survival, as compared to a MazEF-strain. This indicates that the effect of MazEF on RNA phage infection dynamics occurs through direct interference with phage replication and lysis, instead of an indirect effect mediated through abortive infection.

**Figure 2.**
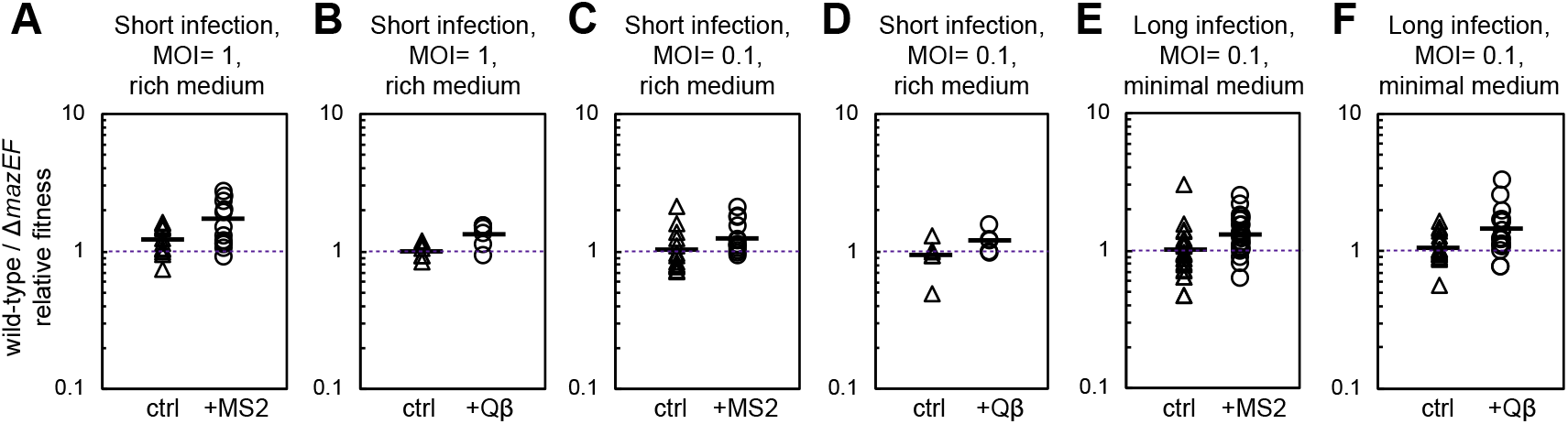
Competition assays between the *E. coli* wild-type and Δ*mazEF* strains during challenge with RNA phage. Exponentially growing *E. coli* wild-type and Δ*mazEF* strains harboring F plasmid (∼10^7^ cells/ml) were initially mixed approximately 1:1; one part of the mixed culture served as a control (triangles) while the other part of the mixed culture was challenged with RNA phage (circles). Phage **A)** MS2 (12 independent replicates, *t*-test *P*= 0.005), or **B)** Qβ (6 replicates, *P*= 0.03) was added at MOI= 1, and phage **C)** MS2 (13 replicates, *P*= 0.03), or **D)** Qβ (6 replicates, *P*= 0.04) was added at MOI= 0.1. Infections were carried out for *t*= 90 min for MS2 and *t*= 105 min for Qβ in rich LB medium. Phage **E)** MS2 (25 replicates, *P*= 0.0004), or **F)** Qβ (18 replicates, *P*= 0.01) was added at MOI= 0.1, and the cultures were exposed to RNA phage overnight in minimal medium. The Wrightian fitness of the wild-type strain relative to the fitness of the Δ*mazEF* strain at time *t*, was calculated as W= (CFUwt_t_/CFUwt_0_) / (CFUΔ*mazEF*_t_/CFUΔ*mazEF*_0_), where CFUwt_0_ and CFUwt_t_ is the density of CFUs for the wild-type strain at time 0 and *t*, respectively and CFUΔ*mazEF*_0_ and CFUΔ*mazEF*_t_ is the density of CFUs at corresponding times for the Δ*mazEF* strain. Strain labeling is described in details in **Material and Methods**. The reciprocal strain labeling was used for overnight infections in minimal medium (linear regression model; MS2 infection significantly affects the relative fitness, *P*= 0.03; Qβ infection significantly affects the relative fitness, *P*= 0.004; strain labeling does not affect the relative fitness, *P*= 0.58).

### MazF induction increases bacterial immunity to RNA phage infection

We hypothesized that the level of *mazF* induction determines the extent of bacterial protection against phage killing. To test this hypothesis, we employed a synthetic system for overexpressing *mazF* from an arabinose-inducible promoter. Previous studies have shown that *mazF* overexpression degrades RNA molecules [Mets et al 2017, Culviner and Laub 2018, Mets et al 2019], and substantially reduces bacterial growth and colony formation [Nikolic et al 2018, Nikolic 2019]. The results of our experiments showed that, while MS2 (Qβ) infection decreased the number of CFUs of uninduced cultures by 99% (87%) on average, it decreased the number of CFUs of *mazF*-induced cultures by only 36% (44%) on average (**Figure 3A-B**). In other words, *mazF*-induced bacterial cultures were less affected by RNA phage infection than uninduced cultures. Higher and longer *mazF* induction resulted in weaker production of RNA phage progeny (**Figure 3C**). It should be noted that the overexpression of *mazF* also had a comparable effect on the progeny production of the obligatory lytic variant of DNA phage λ. This corroborates the results of previous studies on interactions between MazEF and DNA phages [Hazan and Engelberg-Kulka 2004, Engelberg-Kulka and Kumar 2015, Alawneh et al 2016] and suggests that MazEF could contribute to bacterial immunity against both, DNA and RNA phages.

**Figure 3.**
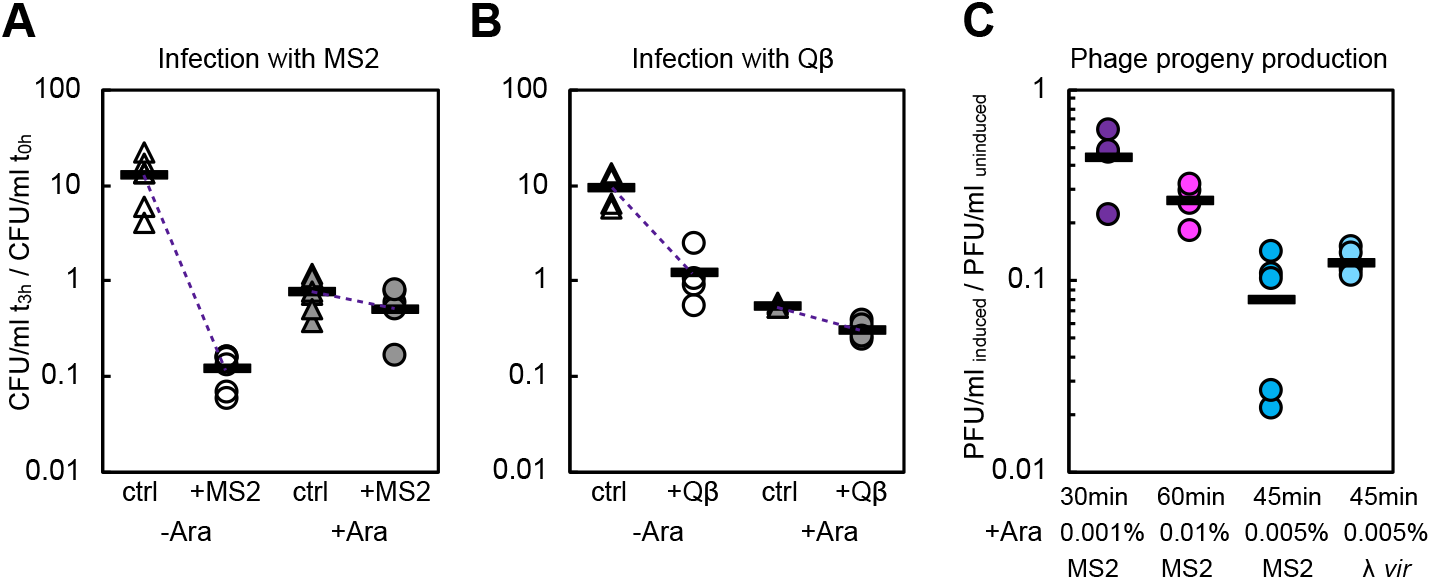
Effect of *mazF* overexpression on formation of bacterial colonies and phage plaques. **A)** Change in the density of CFUs of the Δ*mazEF* F+ strain harboring pBAD-*mazF* was measured for uninfected cultures (triangles) and cultures challenged with RNA phage (circles), with uninduced (open markers) or induced (filled markers) *mazF* expression. The expression of *mazF* was induced with 0.01% arabinose for 30 min prior to addition of the phage. The density of CFUs 3 h after phage addition is reported relative to the density of CFUs at time 0 (∼10^7^ for all replicates). The phage was added at MOI= 0.1, in 6 independent replicate experiments with adding phage MS2, and **B)** 4 replicate experiments with adding phage Qβ. **C)** Decrease in the density of plaque-forming units (PFUs) on bacterial lawns of the Δ*mazEF* F+ pBAD-*mazF* cultures was dependent on the level and time of *mazF* induction. Phage MS2 was used to infect bacterial cultures after inducing *mazF* overexpression with 0.001% Ara for 30 min (purple, 4 replicates), 0.01% Ara for 60 min (magenta, 4 replicates), or 0.005% Ara for 45 min (blue, 5 replicates). In addition, an obligatory lytic variant of phage λ (λ *vir*) was used to infect cultures after inducing *mazF* overexpression with 0.005% Ara for 45 min (light blue, 5 replicates). The density of PFUs for the given phage and induction regime was normalized by the density of PFUs formed on bacterial lawns with uninduced *mazF* expression.

### MazEF delays the time to lysis and increases the size of individual bacterial cells challenged with RNA phage

We next investigated how the presence of the *mazEF* locus affects the progression of phage infection at the single-cell level. We employed a broadly used microfluidic device coupled to a fluorescence microscope [Wang et al 2010], to determine the fate of individual *E. coli* cells upon RNA phage infection. Such single-cell analysis allowed us to better understand bacterial behavior that was not apparent at the population level. *E. coli* cells constitutively expressing *mCherry* fluorescent reporter gene were grown in growth-channels closed at one end and opened to a wider feed-channel. In our analysis, we focused only on the cells residing at the closed end of each microfluidic channel (**Figure 4A**). This microfluidic setup enabled a fast and controlled switch from the phage-free growth medium to the growth medium containing MS2 or Qβ phage lysate. Once the RNA phage were introduced, phage induced-bacteriolysis was detected as the loss of mCherry fluorescence (**Figure 4B**).

**Figure 4.**
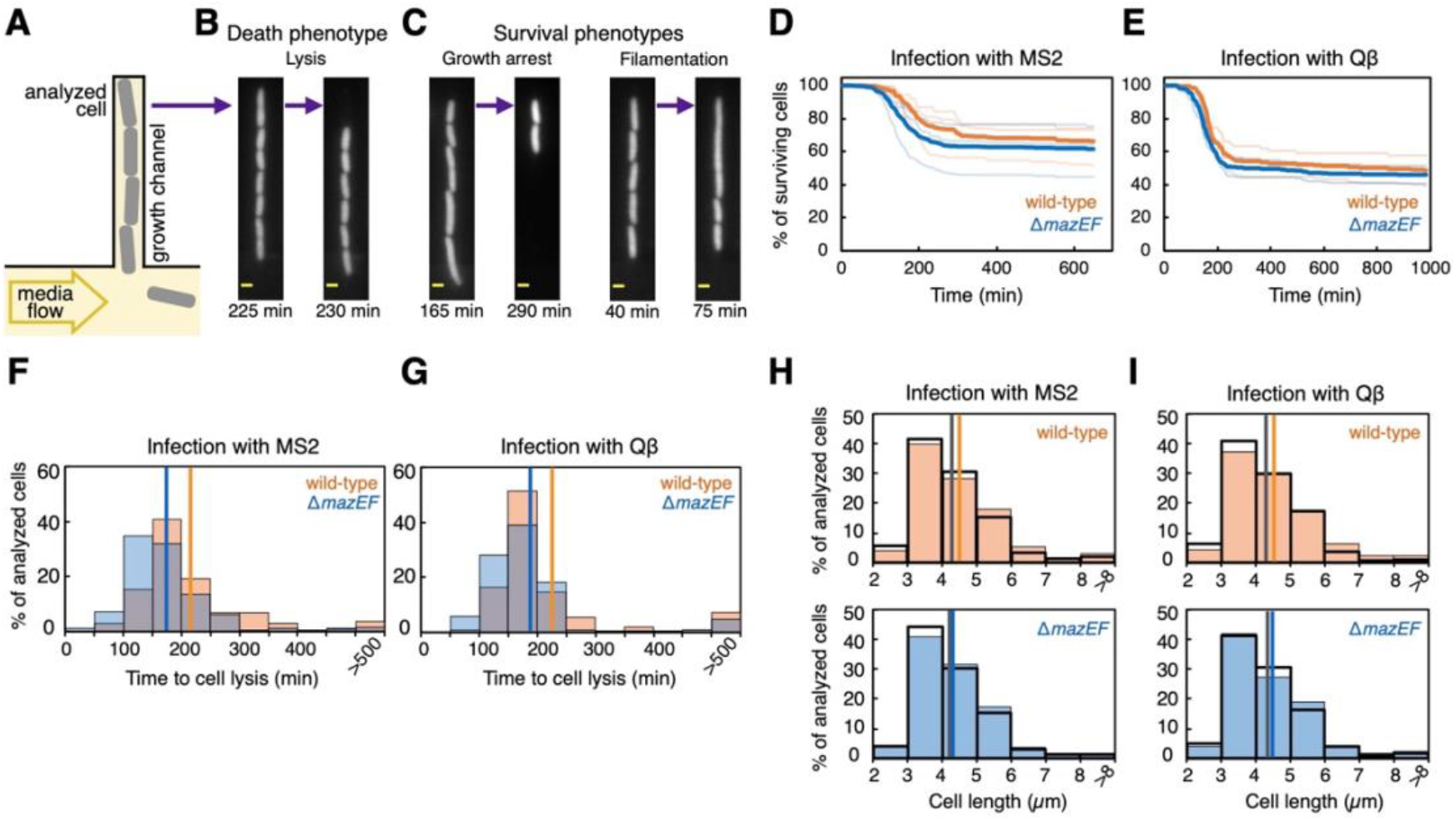
Single-cell analysis of RNA phage infection. **A)** Schematic of the microfluidic device used. *E. coli* cultures were grown for 3 h in the microfluidic device to allow for steady-state growth. Bacteria were then subjected to media containing MS2 or Qβ phage at the density of 10^9^ particles/ml. **B)** Fluorescence microscopy images show growth channels populated with *E. coli* wild-type cells exposed to the medium containing phage MS2. Yellow scale bar represents 1 μm. Lysis of bacterial cells was detected as loss of mCherry fluorescence. **C)** The fraction of cells that did not lyse when challenged with RNA phage exhibited three phenotypes: elongation and division, growth arrest, and elongation and filament formation. **D)** The plots show the percentage of surviving cells as a function of time (only the cells residing at the end of each growth-channel were analyzed during time-lapse experiments). By the end of the experiment, phage MS2 induced lysis in 35.6% of the total 289 wild-type cells (orange) and in 40.9% of the total 428 Δ*mazEF* cells (blue), 3 experiments for each strain. Brighter lines represent measurements of individual experiments, darker lines are the mean values for each strain. **E)** Phage Qβ induced lysis in 54.3% of the total 199 wild-type cells and in 55.3% of the total 329 Δ*mazEF* cells, 2 experiments for each strain. **F)** The histograms show distribution of times to lysis for individual cells, i.e. the duration from switching to the phage-containing media, to cell lysis. 103 *E. coli* wild-type cells lysed on average 213 min after being challenged with MS2 (orange bins; the mean value indicated with orange line), while 175 *E. coli* Δ*mazEF* cells (blue bins; the mean value indicated with blue line) lysed on average after 173 min (Mann-Whitney test *P*= 1.6e-05), with overlapping parts in the histograms depicted in grey. **G)** The analyzed 108 *E. coli* wild-type cells lysed on average 226 min after being challenged with Qβ, while 182 *E. coli* Δ*mazEF* cells lysed on average after 189 min (*P*= 0.004). **H)** The histograms show distributions of cell lengths 5 min before adding phage (white bins with black lines), and 200 min after adding RNA phage (orange bins for the wild-type cells, and blue bins for the Δ*mazEF* cells). We analyzed all cells in static images corresponding to the indicated time points. The length of wild-type cells increased significantly from 4.34 μm (grey line) to 4.51 μm (orange line) on average during MS2 exposure (1644 wild-type cells before and 1521 cells after adding MS2, *P*= 0.004), while the increase in the length of Δ*mazEF* cells from 4.31 μm (grey line) to 4.36 μm (blue line) on average was not significant (2434 wild-type cells before and 1907 cells after adding MS2, *P*=0.054). There were no differences in the length of wild-type and Δ*mazEF* cells prior MS2 exposure (*P*= 0.65). **I)** During Qβ exposure, the length of wild-type cells increased significantly from 4.29 μm to 4.57 μm on average (1168 wild-type cells before and 909 cells after adding Qβ, *P*= 0.0001), while the increase in the length of Δ*mazEF* cells from 4.34 μm to 4.45 μm on average was not significant (1793 wild-type cells before and 1231 cells after adding Qβ, *P*= 0.13). There were no differences in the length of wild-type and Δ*mazEF* cells prior Qβ phage exposure (*P*= 0.58).

We analyzed growth and lysis of bacterial cells during 650 min of exposure to phage MS2 (**Supplementary Movie S1, Supplementary Movie S2**) and 985 min of exposure to phage Qβ. Our data indicated three main growth phenotypes during the periods of RNA phage exposure. First, we observed bacterial cells that continued elongating and dividing in the presence of RNA phage. Second, a fraction of bacterial cells displayed growth arrest in response to RNA phage treatment (**Figure 4C**). Third, a fraction of bacterial cells elongated during RNA phage infection and even produced filaments, which is considered a general survival strategy in bacteria [Justice et al 2008]. These findings corroborate a recent study, which has shown that *E. coli* can exhibit the same three survival phenotypes upon DNA phage T4 infection [Attrill et al 2021].

To study the effect of MazEF on RNA phage infection at the level of single cells, we quantified the fractions of *E. coli* wild-type and Δ*mazEF* cells that survived, i.e. did not lyse, after being challenged with RNA phage (**Figure 4D-E**). The absence of *mazEF* increased the rate of death by 1.15- and 1.04-fold following the period of exposure to phage MS2 and Qβ, respectively. In general, we observed that not all cells lysed when challenged with RNA phage, even if they did not carry the *mazEF* locus. First, it has been previously reported that a fraction of the *E. coli* population might not produce F-pili at all, and that the number of F-pili per cell, as well the length of F-pili, may vary substantially between genetically identical *E. coli* F+ cells [Biebricher and Dueker 1984]. Second, RNA phage infection can promote detachment of F-pili early during the infection process, and block further synthesis of F-pili, both mechanisms rendering bacterial cells less susceptible to subsequent infection [Harb et al 2020]. Third, F-pili may not be entirely accessible for RNA phages to adsorb to them in a confined environment such as the growth channel of the microfluidic device. Fourth, RNA phages have been vastly understudied until very recently, and currently unknown antiphage mechanisms might still be in play in both the wild-type and the Δ*mazEF* strain [Krishnamurthy et al 2016, Callanan et al 2018, Neri et al 2022].

In addition to the overall probability of cell death, we studied how MazEF affects the time to *E. coli* cell lysis, which is an important parameter of phage-host population dynamics. The presence of the *mazEF* locus significantly prolonged the time to MS2-induced lysis by 23% on average (**Figure 4F)**, and the time to Qβ-induced lysis by 19% in challenged *E. coli* cells (**Figure 4G**). By comparing two types of growth media, we observed that the average time to lysis was dependent on the cultivation medium (**Supplementary Figure S4**). Overall, these results suggest that MazEF acts as the first line of defense against RNA phages, reducing initial damage to the infected bacterial population in two ways – by decreasing the probability of cell lysis and by delaying progression of the RNA phage infection within the population.

Finally, we compared the distribution of cell lengths for *E. coli* wild-type and Δ*mazEF* cells. To this end, we analyzed all cells (including cells other than those at the end of the growth-channels) from static images obtained just before and 200 minutes after being challenged with RNA phage. The length of analyzed wild-type cells increased significantly, by 4% on average after adding phage MS2, while the 1% increase in the average length of Δ*mazEF* cells was not significant (**Figure 4H**). Similarly after adding phage Qβ, the length of wild-type cells increased significantly, by 7% on average, and the 2% increase in the average length of Δ*mazEF* cells was not significant (**Figure 4I**). This analysis suggests that bacterial cells that are challenged with RNA phage are longer than cells growing in non-stressful conditions, and that the cell length increase is facilitated by *mazEF*.

### RNA phage genomes exhibit bias against ACA sequences

To extend our analysis outside of the context of the studied model systems, we investigated the abundance of *E. coli*’s MazF recognition sequence ACA in the genomes of RNA phages. For bacterial sequence-specific defense systems targeting DNA phages, such as restriction-modification and CRISPR-Cas systems, avoidance of recognition sequences in the DNA phage genomes is a frequent phenomenon, most likely resulting from selection imposed by these defense systems [Sharp 1986, Kupczok and Bollback 2014, Pleška et al 2016, Pleška and Guet 2017]. If MazF indeed cleaves RNA phage genomes, we should expect similar signatures of selection against ACA recognition sequences in the RNA phage genomes.

To test whether ACA recognition sequences are indeed avoided in RNA phage genomes, we first analyzed genomes that have been deposited as reference RNA phage genomes in the Viral Genomes Database of the National Center for Biotechnology Information (NCBI). These mostly include phages infecting *Escherichia* species, together with phages infecting *Pseudomonas aeruginosa, Caulobacter crescentus* and *Acinetobacter baumannii* respectively, as well as one phage with the broad host range that includes *Escherichia* and *Pseudomonas* species [Olsen and Shipley 1973] (**Table 1**). *mazF* homologs have been also found in the genomes of *P. aeruginosa* and *A. baumannii*, however their sequence specificity has not been investigated so far [Jurenaite et al 2013, Valadbeigi et al 2017]. Here we analyzed the frequency of ACA sites in the collected genomes, and found that the ACA trinucleotide was less frequent than expected in the RNA phage genomes, i.e. their relative ACA frequency was less than 1 (**Figure 5A**). Phages that do not have ssRNA genomes and ssRNA viruses that infect other hosts than Bacteria did not show underrepresentation of the ACA trinucleotide.

**Table 1.**
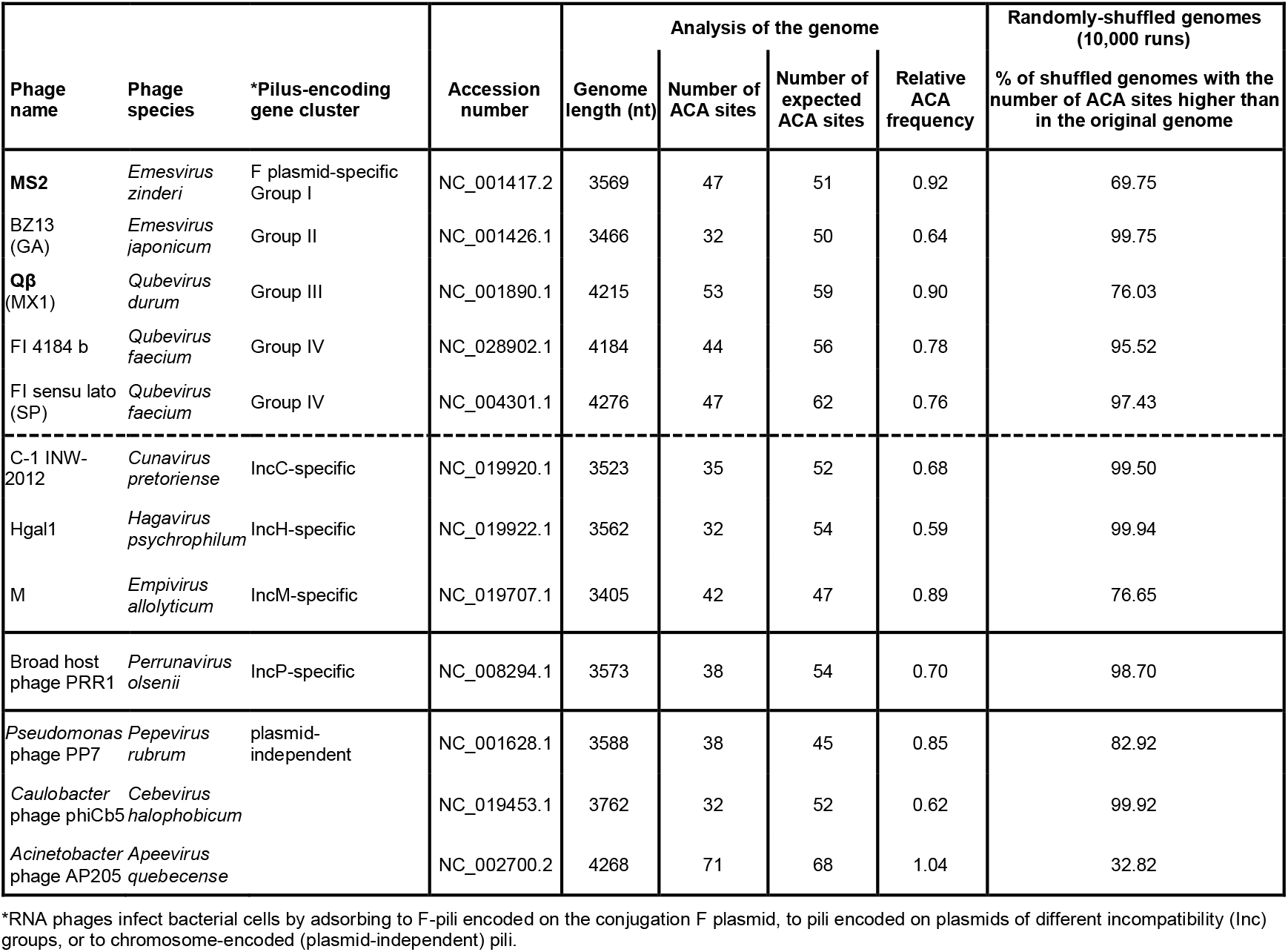
Bioinformatic analysis of RNA phage genomes from the NCBI Reference Sequence Database.

**Figure 5.**
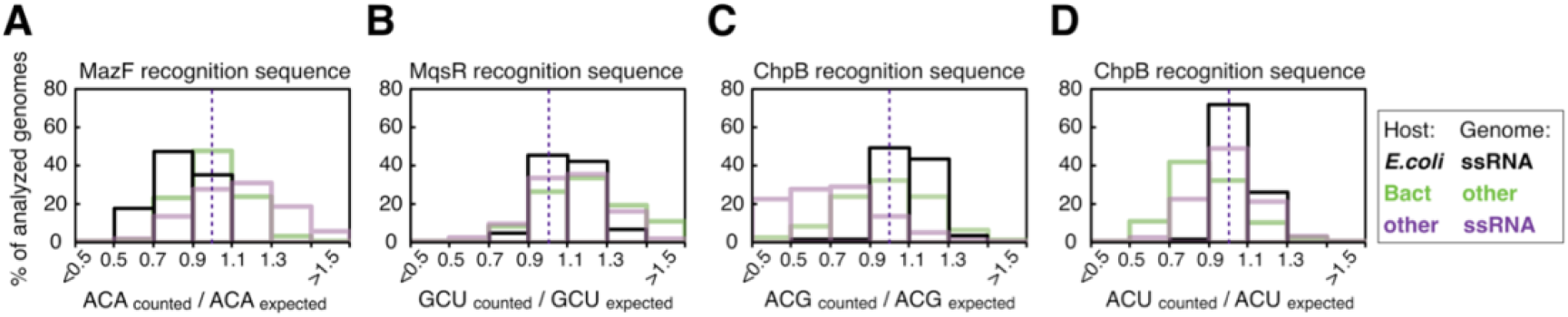
Distributions of relative frequencies for toxin recognition sequences in viral genomes. We analyzed relative frequencies of the recognition sequences of *E. coli*’s toxins MazF, MqsR, and ChpB, specifically **A)** ACA, **B)** GCU (or GCT) **C)** ACG, and **D)** ACU (or ACT) trinucleotides. Relative frequencies were calculated as the actual percentage of the recognition sites relative to their expected percentage in the genome. Black bins show relative frequencies in (+)ssRNA phages infecting *E. coli* (N= 57). Green bins show relative frequencies in phages with genomes other than (+)ssRNA infecting Bacteria (N= 2216). Purple bins show relative frequencies in (+)ssRNA viruses infecting organisms other than bacteria (N= 1835). Dashed lines depict relative frequency of 1. Relative frequencies of ACA, GCU, ACG and ACU trinucleotides in the genomes of RNA phages that infect *E. coli* were 0.83, 1.11, 1.09 and 1.03 on average, respectively.

Specifically, the relative ACA frequency of all 9 reference genomes of the phages that infect *Escherichia* was in the range of 0.59 to 0.92, with the average of 0.76 (**Table 1**). The extended analysis that included RNA phage genomes from the NCBI Nucleotide Database (**Supplementary Table S2**) likewise showed that the ACA trinucleotide in those genomes was underrepresented, with the mean relative frequency of 0.83, averaged over 57 partial and complete RNA genomes of phages that infect *Escherichia* (**Figure 5A**). Finally, we performed analysis of the genomes containing randomly-shuffled nucleotides (10,000 shuffled genomes per one phage genome, **Table 1, Supplementary Table S2**). The results showed that the frequency of ACA sites in the artificial genomes with shuffled nucleotides was significantly higher than in the actual genomes of phages that infect *Escherichia*.

Besides MazF, there are two other *E. coli* ribosome-independent sequence-specific type II toxins. MqsR of *E. coli* cleaves RNA at GCU sites, while ChpB cleaves RNA at ACD sites (D is G, A or U, but not C) [Gerdes 2012, Masuda and Inouye 2017]. Even though ChpB is structurally and biochemically similar to MazF, it is a less efficient enzyme than MazF [Gerdes 2012]. Measured relative frequencies of GCU, ACG and ACU trinucleotides were 1.11, 1.09 and 1.03, respectively (**Figure 5B-D**), which showed that unlike ACA, these trinucleotides were not underrepresented in the genomes of RNA phages infecting *E. coli*. This lack of recognition site avoidance could be either due to a lower activity of the corresponding toxins, or various evolutionary constraints that make particular sites difficult to mutate. Taken together, our results suggest that the avoidance of the MazF recognition sequence ACA is a specific genomic signature of RNA phages that predominantly infect *E. coli*.

## DISCUSSION

Most bacterial antiviral mechanisms have been described for protection against phages with double-stranded DNA genomes [Labrie et al 2010]. Several studies have also investigated protection against phages with single-stranded DNA genomes [Samson et al 2013]. In contrast, our knowledge of how bacteria protect themselves against RNA phages relies on only a handful of studies, which are based on either *in vitro* assays, or non-physiological experimental conditions [Klovins et al 1997, Abudayyeh et al 2016, Smargon et al 2017, Strutt et al 2018, Yan et al 2019]. Our results suggest that the MazEF system facilitates protection of *E. coli* against RNA phages. The MazEF system could contribute to *E. coli* protection at two levels: directly, through MazF-mediated cleavage of RNA phage genomes [Zorzini et al 2016], and indirectly, through MazF-mediated reduction in overall bacterial translation and subsequent bacterial growth decline [Nikolic et al 2018], leading to decreased susceptibility to phage infection. The results of our experiments indicate that MazEF does not predominantly stimulate bacterial cell death upon RNA phage infection, as it is the case with TA Abi systems that promote lysis of infected bacterial cells before the phage can complete its replication cycle [Labrie et al 2010]. It is conceivable that the MazEF-mediated bacterial antiphage strategy is to slow down the spread of RNA phage infection within bacterial populations, and delay RNA phage population growth [Abedon 2017]. Finally, it is plausible that TA systems with toxins that degrade RNA without sequence-specificity, can also serve as bacterial defense mechanisms against RNA phages, and that TA systems exhibit higher or lower efficiency against RNA phages, similarly to the various degrees of efficiency of restriction-modification systems against DNA phages [Pleška and Guet 2017].

Phages are considered to be drivers of bacterial evolution. Bacteria have evolved an array of robust strategies, such as restriction-modification and CRISPR-Cas systems, to protect themselves against DNA phage predation [van Houte et al 2016, Negri et al 2021]. TA systems, such as RnlAB, LsoAB, and MazEF can also play a role in defense against DNA phages [Otsuka 2016, Jurenas et al 2022]. It has been recently shown that MazEF interrupts progression of the lytic cycle of a DNA phage infecting *Bacillus subtilis* [Cui et al 2022]. Furthermore, MazEF system of *E. coli* interacts with DNA phages P1 [Hazan and Engelberg-Kulka 2004], λ [Engelberg-Kulka and Kumar 2015] and T4 [Alawneh et al 2016]. As TA systems have already been regarded as the primary reservoirs of defense agents against DNA phages in *E. coli* [Vassallo et al 2022], this study considers TA systems as the primary line of defense against RNA phages. Moreover, phages have also evolved mechanisms to avoid these TA systems-dependent antiphage strategies [Alawneh et al 2016, Otsuka 2016]. Our analysis indicates that ACA sites in the RNA phage genomes are likely selected against, possibly because phages with fewer ACA sites are more likely to evade MazF action. Together, previous and our results suggest that phage-bacteria coevolution can be mediated by TA systems [Otsuka 2016, Jurenas et al 2022].

In 2015, the *Tara* expedition scientists have provided a large-scale study on the influence of DNA viruses on global ecosystems [Brum et al 2015]. However only recently, the *Tara* consortium has recognized the necessity to investigate RNA viruses in-depth [Dominguez-Huerta et al 2022, Zayed et al 2022]. The impact of RNA viruses has therefore started to be acknowledged not only for global health (e.g. SARS coronavirus, Zika virus, common cold rhinovirus) but also in ecology, evolution and applied sciences. The recent literature suggests that our knowledge of the genomic diversity and host range of RNA phages has been vastly underestimated [Krishnamurthy et al 2016, Callanan et al 2018, Neri et al 2022]. Namely, the Taxonomy Report of the International Committee on Taxonomy of Viruses (ICTV) currently lists 882 bacteria-infecting (+)ssRNA phage species, in contrast to only four (+)ssRNA phage species listed in 2020 [Lefkowitz et al 2017, Callanan et al 2021, Walker et al 2022]. Thus, guiding research towards understanding RNA phage biology, identifying bacterial defense mechanisms against RNA phages, and elucidating how RNA phages affect the dynamics of host-associated commensal and pathogenic bacterial populations, could not be more timely.

## DATA AVAILABILITY

All datasets generated in this study are available within the Supplementary Data (‘Supplementary Datasets.zip’).

## AUTHOR CONTRIBUTIONS

NN designed the study; NN, TB, MP and CCG designed the experiments; NN and TB performed the experiments; NN did image and data analysis; NN, MP and CCG interpreted the data; NN, MP and CCG wrote the manuscript with input from TB.

## ACKNOWLEDGEMENTS

The authors are grateful to Kathrin Tomasek, Lisa Butt, Chris Estell, Alys Jepson, Stefano Pagliara, Remy Chait, Steve West, Vicki Gold, Josh Eaton, Ivana Gudelj, and Rob Beardmore for useful discussions and technical support. The authors thank Laurence Van Melderen for sharing the strains. We acknowledge the IST Austria Lab Support Facility and LSI Technical Services Team at the University of Exeter. NN is grateful to Fabrice Gielen for his support.

