## Supplementary Information for "Bacterial toxin-antitoxin system MazEF as a native defense mechanism against RNA phages in *Escherichia coli*"

**Supplementary Figures**
Figures S1-S4

**Supplementary Tables**
Tables S1 and S2

**Supplementary Movies**
Movies S1 and S2

**Supplementary Datasets**
Source data for experimental results and bioinformatic analysis

**Supplementary Methods**

**Supplementary References**

### Supplementary Figures

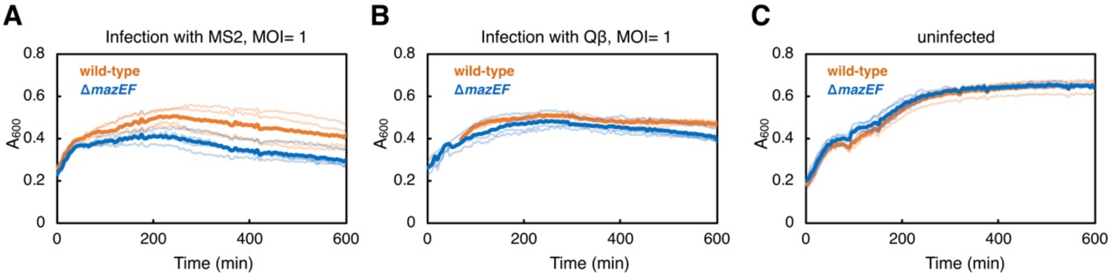


**Supplementary Figure S1.** Bacterial population dynamics during RNA phage infection (a replicate experiment).

Changes in population biomass of the wild-type (orange line) and Δ*mazEF* (blue line) strains were recorded in a plate-reader as $A_{600}$: **A)** after adding phage MS2 at time 0, MOI= 1 (4 replicates, one *t*-test per each time point during period 540-600 min, 13 *t*-tests in total, all *P*< 0.043 except for the time point 555 min with *P*= 0.051), **B)** phage Qβ, MOI= 1 (4 replicates, *t*-tests during period 540-600 min *P*< 0.037), or **C)** without adding phage (4 replicates, *t*-tests during period 540-600 min *P*> 0.72). Brighter lines represent measurements of individual cultures, darker lines are the mean values for each strain. After ten hours of exposure to phage MS2 (Qβ), biomass of the *E. coli* wild-type populations was on average 38% (20%) larger than biomass of the *E. coli* Δ*mazEF* populations.

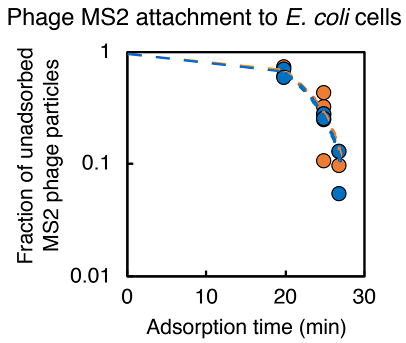


**Supplementary Figure S2.** Attachment of phage MS2 to the *E. coli* wild-type and Δ*mazEF* cells. Between $10^6$-$10^7$ bacterial cells of *E. coli* wild-type (orange circles) and isogenic Δ*mazEF* F+ strains (blue circles) were infected with $10^4$-$10^5$ phage particles of phage MS2 (8 replicates each strain). Bacteria-phage mixtures were incubated at 37°C, without shaking, for the adsorption period of 20 min, 25 min, or 27 min. Afterwards, the samples were plated on phage plates to determine the total phage count ('All'). An aliquot of the mixture was filter-sterilized, and plated on phage plates to determine the fraction of unadsorbed MS2 phage particles ('Unadsorbed'). Detailed experimental protocols are in **Methods** and **Supplementary Methods**. The fraction of unadsorbed MS2 phage particles was determined as $[PFU/ml]_{Unadsorbed}$ / $[PFU/ml]_{All}$. There were no differences in the phage MS2 adsorption between the wild-type and Δ*mazEF* strains.

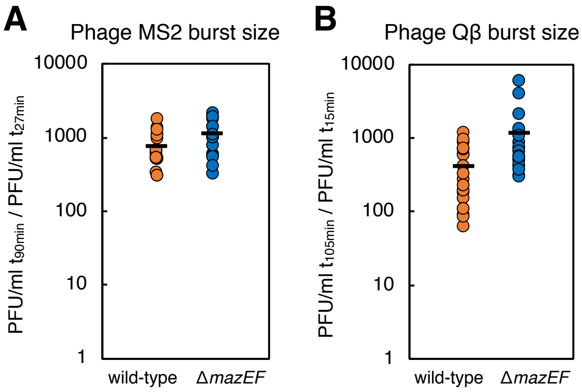


**Supplementary Figure S3.** Burst size of RNA phages infecting *E. coli* wild-type and Δ*mazEF* strains.

Between $10^6$-$10^7$ bacterial cells of *E. coli* wild-type (orange circles) and isogenic Δ*mazEF* strains (blue circles) harboring F plasmid were infected with $10^4$-$10^5$ particles of: **A)** phage MS2 (17 independent replicates) or **B)** phage Qβ (19 replicates). Burst size was calculated as the ratio of the number of PFUs (plaque forming units) at the end of one infection cycle at time point $t_2$ to the initial number of PFUs measured at time point $t_1$ just after phage adsorption. For MS2 experiments, $t_1$= 27 min and $t_2$= 90 min. For Qβ experiments, $t_1$= 15 min and $t_2$= 105 min. Some experiments slightly deviated from these infection protocols, as described in details in **Supplementary Methods**. The plots show that significantly fewer plaques were produced on the wild-type than on the Δ*mazEF* cultures after MS2 infection (linear regression model; the strain genotype significantly affects MS2 plaque formation, *P*= 0.041; the infection protocols do not affect MS2 plaque count, *P*= 0.89, 0.81 and 0.21), as well as after Qβ infection (linear regression model; the strain genotype significantly affects Qβ plaque formation, *P*= 0.047; the infection protocols do not affect Qβ plaque count, *P*= 0.45 and 0.21).

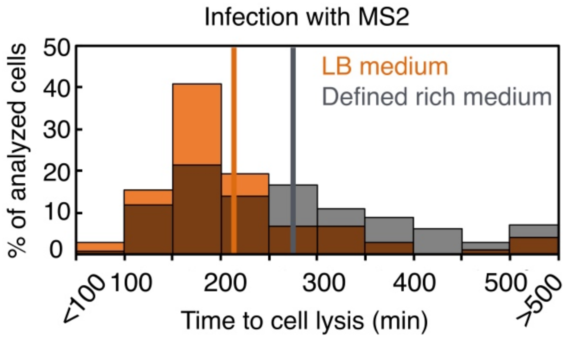


**Supplementary Figure S4.** Time to bacteriolysis in a microfluidic device depends on the cultivation medium.

We investigated how pre-culturing conditions affect the response of bacterial cells to phage MS2. To this end, *E. coli* wild-type strain NN242-cat F+ pre-cultivated either in LB medium or in defined rich medium containing 0.5% casamino acids and 0.2% glucose, was loaded into a microfluidic device. *E. coli* cells were grown in LB medium in the microfluidic device for at least 3 h before the medium was switched to the same medium, but containing $10^9$ MS2 phage particles/ml. The cell fate was monitored for 650 min after introducing the phage. Phage MS2 induced lysis in 35.6% of a total of 289 cells pre-cultivated in LB medium (3 experiments, the same dataset as in **Figure 4**), and in 36.9% of 393 cells pre-cultivated in defined rich medium (2 experiments). Lysis of the 103 *E. coli* cells pre-cultivated in LB medium occurred on average 213 min after the addition of MS2 (orange bins; the mean value indicated with orange line), while lysis of the 145 cells pre-cultivated in defined rich medium occurred on average 277 min after the addition of MS2 (grey bins; the mean value indicated with dark grey line), i.e. the average time to lysis was 30% longer (Mann-Whitney test, *P*= 3.6e-06).

### Supplementary Tables

**Supplementary Table S1**. Phages, bacterial strains and plasmids used in this study.

| Name | Relevant characteristics | Source or reference |
|---|---|---|
| *Enterobacteria* phage MS2 | *Emesvirus zinderi* (+)ssRNA bacteriophage | DSM 13767, Leibniz Institute DSMZ-German Collection of Microorganisms and Cell Cultures |
| *Enterobacteria* phage Qβ | *Qubevirus durum* (+)ssRNA bacteriophage | DSM 13768, Leibniz Institute DSMZ-German Collection of Microorganisms and Cell Cultures |
| *Escherichia* phage λ *vir* | An obligatory lytic variant of phage *Escherichia virus Lambda* dsDNA bacteriophage | Guet Collection |
| MG1655 | *E. coli* K-12 wild-type F-, λ-, *ilvG-, rfb-50, rph-1* | [Blattner et al 1997] Bergmiller Strain Collection |
| MG1655 LVM | *E. coli* K-12 wild-type from LVM Lab | [Tsilibaris et al 2007] Laurence Van Melderen Strain Collection |
| W1485 | *E. coli* Lederberg W1485; F+ *met str*, T1s T6s λ- Classical F+ donor in conjugation | DSM 5695, Leibniz Institute DSMZ-German Collection of Microorganisms and Cell Cultures |
| Hfr 3000 U 432 | *E. coli* Hfr 3000 U 432 | DSM 5210, Leibniz Institute DSMZ-German Collection of Microorganisms and Cell Cultures |
| LVM101 | MG1655 Δ*mazEF* | [Tsilibaris et al 2007] Laurence Van Melderen Strain Collection |
| TB315 | MG1655 Δ*araA*::*kanR* | Bergmiller Strain Collection |
| TB194 | MG1655 *att*P21::λ$P_R$-*mCherry*::frt-*cat* constitutive *mCherry* expression | Bergmiller Strain Collection |
| NN238 | LVM101 Δ*araA*::*kanR* Δ*araA*::*kanR* P1-transduced from TB315 | This study |
| NN239 | LVM MG1655 *att*HK022::frt-*cat* CRIM protocol [Haldimann & Wanner 2001] | This study |
| NN241 | LVM101 *att*HK022::frt-*cat* CRIM protocol [Haldimann & Wanner 2001] | This study |
| NN242-cat | LVM MG1655 *att*P21::λ$P_R$-*mCherry*::frt-*cat* *att*P21::λ$P_R$-*mCherry*::frt-*cat* P1-transduced from TB194 | This study |
| NN243-cat | LVM101 *att*P21::λ$P_R$-*mCherry*::frt-*cat* *att*P21::λ$P_R$-*mCherry*::frt-*cat* P1-transduced from TB194 | This study |
| NN242 | LVM MG1655 *att*P21::λ$P_R$-*mCherry*::frt *cat*-cassette removed with Flp recombinase [Cherepanov and Wackernagel 1995] | This study |
| NN243 | LVM101 *att*P21::λ$P_R$-*mCherry*::frt *cat*-cassette removed with Flp recombinase [Cherepanov and Wackernagel 1995] | This study |
| TB315 F+ | F plasmid from W1485 introduced by conjugation in TB315 | This study |
| NN238 F+ | F plasmid from W1485 introduced by conjugation in NN238 | This study |
| NN239 F+ | F plasmid from W1485 introduced by conjugation in NN239 | This study |
| NN241 F+ | F plasmid from W1485 introduced by conjugation in NN241 | This study |
| NN242-cat F+ | F plasmid from W1485 introduced by conjugation in NN242-cat | This study |
| NN243-cat F+ | F plasmid from W1485 introduced by conjugation in NN243-cat | This study |
| pAH68-frt-cat | pAH68 Δ*bla*::frt-*cat* | Bergmiller Strain Collection |
| pAH69 | helper plasmid for integration in *att*HK022 | [Haldimann and Wanner 2001] |
| pBAD-*mazF* | CamR, p15A ori, pBAD33 backbone, $P_{BAD}$-*mazF* Ara-inducible *mazF* expression | [Amitai et al 2004] |

**Supplementary Table S2.** Extended bioinformatic analysis of RNA phage genomes. Analysis of sequences retrieved from the NCBI Nucleotides Database, comprising 28 genomes from [Friedman et al 2009], 20 genomes from [Krishnamurthy et al 2016] as well as complete and partial genomes of (+)ssRNA phages with the length of more than 1000 nucleotides and excluding genomes obtained from experimentally evolved strains (81 genomes in total, on average 1.24% of ACA sites per genome). Ten phage genomes exhibited the fraction of ACA trinucleotides higher than the expected fraction of ACA sites (highlighted in purple). These include phage isolates with the unknown host range [Krishnamurthy et al 2016], *Escherichia* phage M11 [Beekwilder et al 1995], and *Acinetobacter* phage AP205.

| Species | Genome length (nt) | Number of ACA sites | ACA sites (%) | Expected ACA (%) | Relative ACA frequency | % of shuffled genomes with the number of ACA sites higher than in the original genome |
|---|---|---|---|---|---|---|
| 'KT462702.1 Leviviridae sp. isolate AVE008 hypothetical protein and replicase genes, partial cds' | 2565 | 62 | 2.42 | 1.76 | 1.38 | 0.5 |
| 'KT462712.1 Leviviridae sp. isolate ESE003 hypothetical protein gene, complete cds; and replicase gene, partial cds' | 1366 | 25 | 1.83 | 1.45 | 1.26 | 8.47 |
| 'KT462707.1 Leviviridae sp. isolate AVE013 replicase gene, partial cds' | 946 | 15 | 1.59 | 1.35 | 1.18 | 19.68 |
| 'KT462704.1 Leviviridae sp. isolate AVE010 hypothetical protein gene, partial cds; maturation gene, complete cds; and hypothetical protein gene, partial cds' | 1753 | 34 | 1.94 | 1.74 | 1.12 | 21.64 |
| 'KT462701.1 Leviviridae sp. isolate AVE007 maturation gene, partial cds' | 3580 | 54 | 1.51 | 1.41 | 1.07 | 26.1 |
| 'AF052431.1 Bacteriophage M11 A-protein, coat protein, A1-protein, and replicase genes, complete cds' | 4217 | 60 | 1.42 | 1.33 | 1.07 | 26.3 |
| 'KT462694.1 Leviviridae sp. isolate AVE000 hypothetical protein gene, partial cds; and hypothetical protein, maturation, hypothetical protein, and replicase genes, complete cds' | 4977 | 90 | 1.81 | 1.71 | 1.06 | 26.36 |
| 'KT462709.1 Leviviridae sp. isolate ESE000 maturation and replicase genes, partial cds' | 3426 | 56 | 1.64 | 1.53 | 1.07 | 26.78 |
| 'NC_002700.2 Acinetobacter phage AP205, complete genome' | 4268 | 71 | 1.66 | 1.60 | 1.04 | 33.93 |
| 'KT462705.1 Leviviridae sp. isolate AVE011 maturation and hypothetical protein genes, partial cds' | 1422 | 26 | 1.83 | 1.82 | 1.01 | 43.36 |
| 'EU403427.1 Enterobacteria phage FI isolate HB-P22 replicase gene, partial cds' | 1159 | 15 | 1.30 | 1.35 | 0.96 | 49.68 |
| 'FJ483843.1 Enterobacteria phage Qbeta strain VK, complete genome' | 4218 | 55 | 1.30 | 1.33 | 0.98 | 52.49 |
| 'KT462700.1 Leviviridae sp. isolate AVE006 maturation gene, partial cds; and hypothetical protein and replicase genes, complete cds' | 3609 | 59 | 1.64 | 1.67 | 0.98 | 53.44 |

| | | | | | | |
|---|---|---|---|---|---|---|
| 'GQ153931.1 Enterobacteria phage Qbeta isolate QB_ancestral A2 maturation protein, A1 read-through protein, coat protein, and replicase genes, complete cds' | 4198 | 51 | 1.22 | 1.25 | 0.97 | 54.23 |
| 'AB971354.1 Enterobacteria phage Qbeta RNA, complete genome, isolate: Anc(P1)' | 4217 | 51 | 1.21 | 1.25 | 0.97 | 54.47 |
| 'FJ483840.1 Enterobacteria phage Qbeta strain TW18, complete genome' | 4218 | 53 | 1.26 | 1.29 | 0.97 | 54.99 |
| 'JQ966308.1 Enterobacterio phage MS2 isolate DL54, partial genome' | 3398 | 47 | 1.38 | 1.45 | 0.95 | 59.89 |
| 'AY099114.1 Bacteriophage Qbeta A2 maturation protein, coat protein, A1 read-through protein, and Q beta replicase genes, complete cds' | 4160 | 49 | 1.18 | 1.24 | 0.95 | 61.26 |
| 'MK213795.1 Escherichia phage MS2, complete genome' | 3526 | 48 | 1.36 | 1.43 | 0.95 | 62.19 |
| 'KT462703.1 Leviviridae sp. isolate AVE009 maturation gene, partial cds' | 2094 | 30 | 1.43 | 1.54 | 0.93 | 62.46 |
| 'EU403428.1 Enterobacteria phage FI isolate HB-P24 replicase gene, partial cds' | 1159 | 12 | 1.04 | 1.20 | 0.87 | 63.24 |
| 'FJ799467.1 Enterobacterio phage MS2 clone MS2anc assembly protein (MS2g1), coat protein (MS2g2), lysis protein (MS2g3), and replicase protein (MS2g4) genes, complete cds' | 3481 | 47 | 1.35 | 1.45 | 0.93 | 65.34 |
| 'KT462698.1 Leviviridae sp. isolate AVE004 maturation gene, partial cds; and hypothetical protein and replicase genes, complete cds' | 3776 | 65 | 1.72 | 1.83 | 0.94 | 66.32 |
| 'FJ483841.1 Enterobacteria phage Qbeta strain HL4-9, complete genome' | 4221 | 51 | 1.21 | 1.30 | 0.93 | 67.21 |
| 'M31635.1 Bacteriophage fr maturation and coat protein genes, complete cds, and replicase gene, 5'' end' | 1788 | 25 | 1.40 | 1.56 | 0.90 | 67.5 |
| 'X14764.1 Bacteriophage Q-beta mRNA for replicase' | 1964 | 22 | 1.12 | 1.26 | 0.89 | 67.79 |
| NC_001417.2 Enterobacterio phage MS2, complete genome' | 3569 | 47 | 1.32 | 1.43 | 0.92 | 69.27 |
| 'GQ153927.1 Enterobacterio phage MS2 isolate MS2_ancestral assembly protein, coat protein, lysis protein, and replicase genes, complete cds' | 3569 | 47 | 1.32 | 1.43 | 0.92 | 69.45 |
| 'pdbl5TC1IR Chain R, phage MS2 genome' | 3569 | 47 | 1.32 | 1.43 | 0.92 | 69.89 |
| 'EF204940.1 Enterobacteria phage MS2 isolate ST4, complete genome' | 3569 | 46 | 1.29 | 1.41 | 0.92 | 70.9 |
| 'FJ539132.1 Enterobacteria phage FI strain HB-P22, complete genome' | 4241 | 54 | 1.27 | 1.38 | 0.92 | 71.22 |
| 'MH108093.1 Escherichia virus Qbeta isolate Qbeta(wt), partial genome' | 3900 | 44 | 1.13 | 1.24 | 0.91 | 71.71 |
| 'FJ483844.1 Enterobacteria phage Qbeta strain BZ1, complete genome' | 4219 | 51 | 1.21 | 1.32 | 0.91 | 72.8 |
| 'NC_001890.1 Enterobacteria phage Qbeta, complete genome' | 4215 | 53 | 1.26 | 1.39 | 0.90 | 75.18 |
| 'AF059242.1 Bacteriophage MX1, complete genome' | 4215 | 53 | 1.26 | 1.39 | 0.90 | 75.89 |

| | | | | | | |
|---|---|---|---|---|---|---|
| 'AH009279.2 Enterobacteria phage JP501 maturation protein, coat protein, and lysis protein genes, complete cds; and RNA replicase beta chain genes, partial cds' | 2729 | 29 | 1.06 | 1.23 | 0.87 | 75.98 |
| 'FJ483838.1 Enterobacteria phage BZ13 strain T72, complete genome' | 3393 | 46 | 1.36 | 1.51 | 0.90 | 76.28 |
| 'KT462699.1 Leviviridae sp. isolate AVE005 maturation gene, partial cds; and hypothetical protein and replicase genes, complete cds' | 3709 | 52 | 1.40 | 1.56 | 0.90 | 76.65 |
| 'NC_019707.1 Enterobacteria phage M, complete genome' | 3405 | 42 | 1.23 | 1.39 | 0.89 | 76.93 |
| 'AB634835.1 Enterobacteria phage FI genomic RNA, RNA-directed RNA polymerase beta chain region, isolate: 110112RS392' | 1078 | 11 | 1.02 | 1.33 | 0.77 | 78 |
| 'AB627075.1 Enterobacteria phage FI genomic RNA, RNA-directed RNA polymerase beta chain region, isolate: 110112RS395' | 1069 | 12 | 1.12 | 1.45 | 0.78 | 78.52 |
| 'JQ966307.1 Enterobacterio phage MS2 isolate DL52, complete genome' | 3525 | 44 | 1.25 | 1.42 | 0.88 | 79.5 |
| 'KT462706.1 Leviviridae sp. isolate AVE012 hypothetical protein and maturation genes, partial cds' | 1053 | 13 | 1.24 | 1.60 | 0.77 | 80.09 |
| 'EF107159.1 Enterobacteria phage MS2 isolate DL1 assembly protein, coat protein, lysis protein, and replicase genes, complete cds' | 3570 | 43 | 1.21 | 1.39 | 0.87 | 81.32 |
| 'FJ483842.1 Enterobacteria phage Qbeta strain BR12, complete genome' | 4218 | 48 | 1.14 | 1.30 | 0.87 | 82.52 |
| 'LT898437.1 Escherichia virus MS2 isolate EnteroPH_GER_L00928-K20_14-03_2014 genome assembly, complete genome: monopartite' | 3534 | 43 | 1.22 | 1.41 | 0.86 | 82.64 |
| 'NC_001628.1 Pseudomonas phage PP7, complete genome' | 3588 | 38 | 1.06 | 1.24 | 0.85 | 83.33 |
| 'EF204939.1 Enterobacteria phage MS2 isolate J20, complete genome' | 3569 | 42 | 1.18 | 1.38 | 0.85 | 84.8 |
| 'AF195778.1 Enterobacteriophage M12 maturation protein, coat protein, and lysis protein genes, complete cds; and RNA replicase beta chain gene, partial cds' | 3340 | 41 | 1.23 | 1.44 | 0.85 | 85.18 |
| 'AB634836.1 Enterobacteria phage FI genomic RNA, RNA-directed RNA polymerase beta chain region, isolate: 110112RS394' | 1092 | 11 | 1.01 | 1.42 | 0.71 | 85.41 |
| 'KT462711.1 Leviviridae sp. isolate ESE002 hypothetical protein gene, partial cds; and maturation and hypothetical protein genes, complete cds' | 1451 | 18 | 1.24 | 1.61 | 0.77 | 86.43 |
| 'AB624552.1 Enterobacteria phage FI genomic RNA, RNA-directed RNA polymerase beta chain region, isolate: 110112RS397' | 1086 | 10 | 0.92 | 1.35 | 0.68 | 87.73 |
| 'FJ539134.1 Enterobacteria phage FI strain BR1, complete genome' | 4273 | 50 | 1.17 | 1.40 | 0.84 | 89.6 |
| 'X15031.1 Bacteriophage fr RNA genome' | 3575 | 44 | 1.23 | 1.49 | 0.83 | 89.67 |

| | | | | | | |
|---|---|---|---|---|---|---|
| 'KT462708.1 Leviviridae sp. isolate AVE014 replicase gene, partial cds' | 799 | 10 | 1.25 | 1.87 | 0.67 | 89.75 |
| 'EF108465.1 Enterobacteria phage MS2 isolate R17, complete genome' | 3569 | 42 | 1.18 | 1.43 | 0.82 | 90.06 |
| 'M99039.1 Phage Q-beta coat protein and A1 protein genes, complete cds' | 1062 | 9 | 0.85 | 1.32 | 0.64 | 90.14 |
| 'KT462697.1 Leviviridae sp. isolate AVE003 maturation gene, partial cds; and hypothetical protein and replicase genes, complete cds' | 3957 | 49 | 1.24 | 1.49 | 0.83 | 90.46 |
| 'EU403429.1 Enterobacteria phage FI isolate BR1 replicase gene, partial cds' | 1159 | 11 | 0.95 | 1.41 | 0.67 | 90.73 |
| 'KT462710.1 Leviviridae sp. isolate ESE001 maturation gene, complete cds' | 1862 | 28 | 1.51 | 1.93 | 0.78 | 90.99 |
| 'LT821717.1 LeviOr01 virus genome assembly, chromosome: LeviOr01' | 3669 | 38 | 1.04 | 1.29 | 0.80 | 91.29 |
| 'AB624551.1 Enterobacteria phage FI genomic RNA, RNA-directed RNA polymerase beta chain region, isolate: 110112RS393' | 1086 | 10 | 0.92 | 1.44 | 0.64 | 91.97 |
| 'EF108464.1 Enterobacteria phage MS2 isolate DL16, complete genome' | 3569 | 40 | 1.12 | 1.40 | 0.80 | 92.01 |
| 'FJ539133.1 Enterobacteria phage FI strain HB-P24, complete genome' | 4243 | 45 | 1.06 | 1.30 | 0.81 | 92.1 |
| 'AF059243.1 Bacteriophage NL95, complete genome' | 4248 | 46 | 1.08 | 1.36 | 0.80 | 94.25 |
| 'FJ539135.1 Enterobacteria phage FI strain BR8, complete genome' | 4273 | 48 | 1.12 | 1.40 | 0.80 | 94.44 |
| 'KT462696.1 Leviviridae sp. isolate AVE002 maturation, hypothetical protein, and replicase genes, complete cds' | 4220 | 56 | 1.33 | 1.63 | 0.81 | 94.89 |
| 'NC_028902.1 Enterobacteria phage FI 4184 b, complete genome' | 4184 | 44 | 1.05 | 1.34 | 0.78 | 95.48 |
| 'KT462695.1 Leviviridae sp. isolate AVE001 maturation gene, partial cds; and hypothetical protein and replicase genes, complete cds' | 5021 | 56 | 1.12 | 1.39 | 0.80 | 95.68 |
| 'GQ153935.1 Enterobacteria phage SP isolate SP_ancestral maturation protein, read-through protein, coat protein, and replicase genes, complete cds' | 4276 | 48 | 1.12 | 1.43 | 0.79 | 95.79 |
| 'EU403430.1 Enterobacteria phage FI isolate BR8 replicase gene, partial cds' | 1159 | 10 | 0.86 | 1.47 | 0.59 | 95.96 |
| 'NC_004301.1 Bacteriophage SP genomic RNA' | 4276 | 47 | 1.10 | 1.44 | 0.76 | 97.76 |
| 'KT462713.1 Leviviridae sp. isolate ESE004 maturation gene, complete cds' | 1290 | 9 | 0.70 | 1.35 | 0.52 | 98.54 |
| 'AF227250.1 Enterobacteriophage KU1, complete genome' | 3486 | 36 | 1.03 | 1.47 | 0.70 | 98.8 |
| 'NC_008294.1 Pseudomonas phage PRR1, complete genome' | 3573 | 38 | 1.06 | 1.51 | 0.70 | 99 |
| 'FJ483839.1 Enterobacteria phage BZ13 strain DL20, complete genome' | 3458 | 33 | 0.95 | 1.40 | 0.68 | 99.11 |
| 'FJ483837.1 Enterobacteria phage BZ13 strain DL10, complete genome' | 3412 | 33 | 0.97 | 1.43 | 0.68 | 99.17 |
| 'NC_019920.1 Enterobacteria phage C-1 INW-2012, complete sequence' | 3523 | 35 | 0.99 | 1.47 | 0.68 | 99.31 |

| | | | | | | |
|---|---|---|---|---|---|---|
| 'NC_001426.1 Enterobacteria phage GA, complete genome' | 3466 | 32 | 0.92 | 1.43 | 0.64 | 99.65 |
| 'NC_019453.1 Caulobacter phage phiCb5, complete genome' | 3762 | 32 | 0.85 | 1.37 | 0.62 | 99.83 |
| 'NC_019922.1 Enterobacteria phage Hgal1, complete sequence' | 3562 | 32 | 0.90 | 1.51 | 0.59 | 99.97 |

### Supplementary Movies

**Movie S1**. Time-lapse of single cells of the *E. coli* wild-type strain during exposure to phage MS2. (Scale bar corresponds to 2 µm.)
**Movie S2**. Time-lapse of single cells of the *E. coli* Δ*mazEF* strain during exposure to phage MS2.

### Supplementary Methods

**Burst size measurements.** Overnight cultures of the wild-type NN239 F+ or Δ*mazEF* NN241 F+ strains were diluted 1 to 1000 into fresh LB medium and cultivated for 3.5 h. 500 µl of exponentially growing cultures containing $10^6$-$10^7$ bacterial cells were then mixed with 10-100 µl of RNA phage solution containing $10^4$-$10^5$ phage particles. For each replicate experiment, the number of infective centers was determined at time $t_1$ just after phage adsorption, and at time point $t_2$, which corresponds to the length of the phage infection cycle. Just before measuring the number of infective centers at time $t_1$, bacteria-phage samples were washed twice with fresh LB medium to remove all unattached phage particles. Serial dilutions of the samples were made, and 10, 20 or 100 µl of diluted samples were then mixed with 500 µl of bacterial host culture (~$10^6$ cells of strain W1485) in phage soft agar, and plated on phage plates. Standard protocol was $t_1$= 27 min and $t_2$= 90 min for MS2 infection 'regMS', and $t_1$= 15 min and $t_2$= 105 min for Qβ infection 'regQB'. Other protocols for MS2 infection were: 'nowash20' ($t_1$= 20 min, no washing after adsorption), 'nowash25' ($t_1$= 25 min, no washing after adsorption), 'nowash27' ($t_1$= 27 min, no washing after adsorption), as indicated in **Supplementary Datasets**. Other protocols for Qβ infection were: 'ads19' ($t_1$= 19 min), 'ads16tot100' ($t_1$= 16 min, $t_2$= 100 min). Burst size was calculated for each strain as: burst size= $[PFU/ml]_{t2}$ / $[PFU/ml]_{t1}$. For experiments without the washing step after adsorption, we determined the total phage count after adsorption ('All'). An aliquot of the mixture was filter-sterilized through a 0.22 µm-filter, mixed with exponentially growing bacterial cultures of host strain W1485 and soft agar, and plated on phage plates to determine the fraction of unadsorbed MS2 phage particles ('Unadsorbed'). For these experiments, burst size was calculated as: burst size with correction= $[PFU/ml]_{t2}$ / $\{[PFU/ml]_{t1\ All} - [PFU/ml]_{t1\ Unadsorbed}\}$.

**Statistical analysis – linear regression models.** Competition assays with the reciprocal strain labeling were evaluated by the Matlab function *fitlm*, by fitting a linear regression model with the different conditions as categorical predictor variables ('ThePhageType': no phage, MS2 infection, or Qβ infection; 'TypeOfTheLabeling': Type L1= the wild-type strain is Ara+ and the

Δ*mazEF* strain is Ara-, or Type L2= wild-type is Ara- and Δ*mazEF* is Ara+), and relative fitness as the response variable. Likewise, burst size assays were evaluated by the Matlab function *fitlm*, by fitting a linear regression model with the different conditions as categorical predictor variables (‘Strain’: NN239 F+ or NN241 F+; ’Protocol’: different burst size protocols for MS2 or Qβ infection described in the **Supplementary Methods** part **Burst size measurements**), and burst size as the response variable.
